## Supplementary material for "Dissociating task acquisition from expression during learning reveals latent knowledge"

### SUPPLEMENTAL MOVIE LEGENDS

**All supplementary movies are available at circuits.jhu.edu/behaviorvideos**

**Movie S1:** Mouse performing at expert levels in the reinforced context after prolonged training.

**Movie S2:** Video of mouse behaving in the reinforced context (trial block 1500-2000).

**Movie S3:** Video of mouse behaving in the probe context (trial block 1500-2000, immediately after **Movie S2**).

**Movie S4:** Video of mouse behaving in the reinforced context with lever (trial block 1500-2000).

**Movie S5:** Video of mouse behaving in the probe context with (trial block 1500-2000, immediately after **Movie S4**).

SUPPLEMENTAL FIGURES

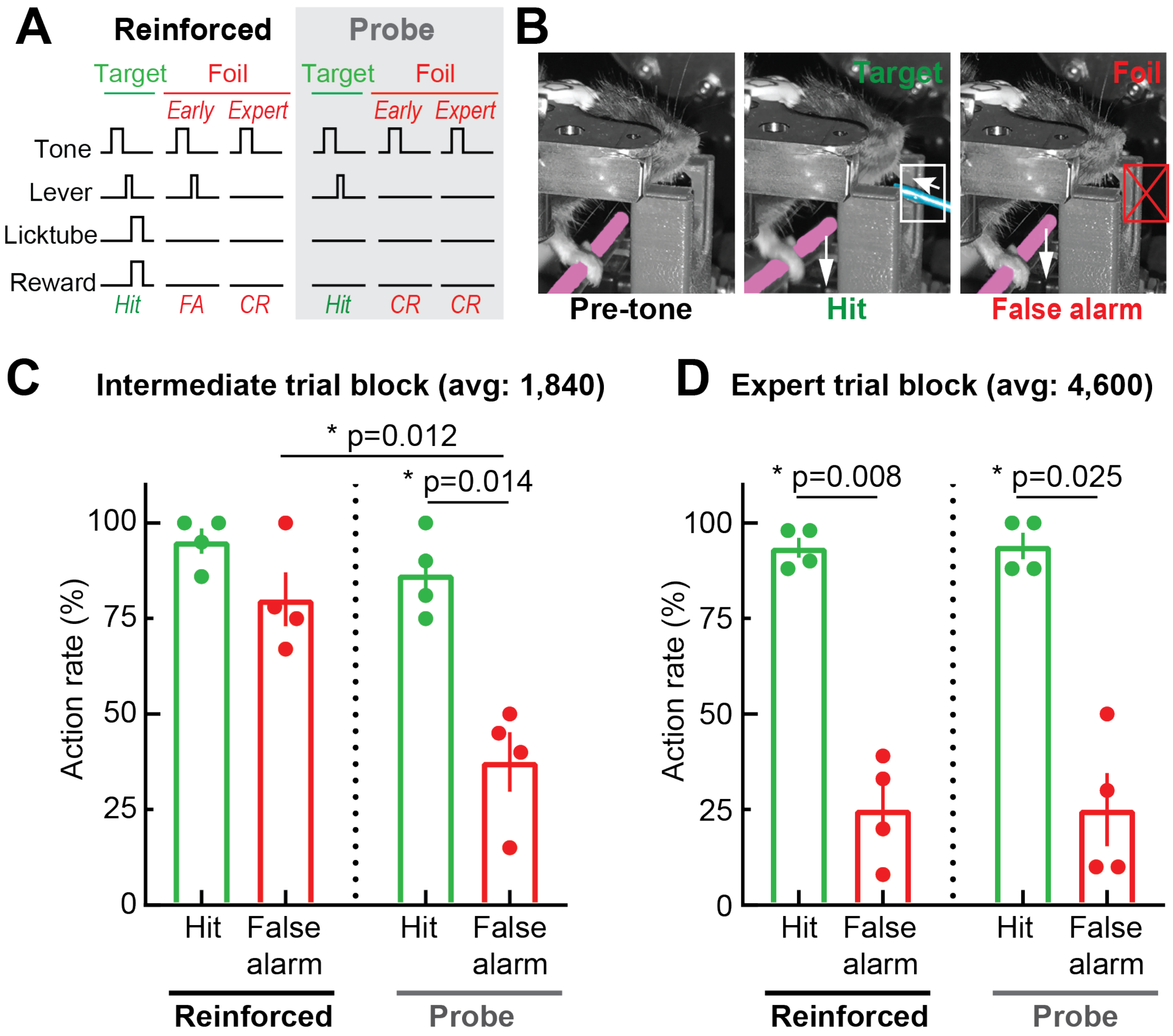


**Figure S1: Lever training dissociates acquisition and expression. A**, Trial structure for lever go/no-go task. In the reinforced context, the licktube was normally absent (low) and was advanced (high) when the mouse correctly pressed the lever to the target tone. In Foil trials, the animal continued to press the lever in response to the tone (false alarm) early in training (trial block 1500-2000) and stopped pressing the lever later in training (>4,000 trials). In the probe context, the animal pressed the lever early in training in response to the target tone but did not do so in response to the foil tone. Note that to transition to the probe context, the animal was given two consecutive target trials; upon pressing the lever, the licktube was not advanced and the animal was then given foil tone trials. **B**, Video frames of the mouse performing the lever task in the reinforced context. Before tone onset, the mouse rested its paw on the lever and the licktube was not present. In a hit trial (target tone), the animal correctly pressed the lever and the licktube delivered a droplet of water. In a false alarm trial (foil tone), the animal incorrectly pressed the lever and the licktube was not advanced. **C**, Quantification of results at an intermediate time point in learning (average trial =1,840; n=4 mice, reinforced context: hit rate=95.3±3.3%, false-alarm rate=80.0±7.1%, p=0.3, probe context: hit rate=86.5±5.4%, false-alarm rate=37.5±7.8%, p=0.014, one-way repeated-measures ANOVA followed by Tukey's post-hoc correction). **D,** Quantification of results at a later time point in learning (average trial=4,600; n=4 mice, reinforced context**:** hit rate=93.5±2.6%, false-alarm rate=25.0±6.9%, p=0.008; probe context**:** hit rate=94.0±3.5%, false-alarm rate=25.0±9.6%, p=0.025, one-way repeated-measures ANOVA followed by Tukey's post-hoc correction). Note that we typically ran "probe" trials only at 2-3 time points with lever training.

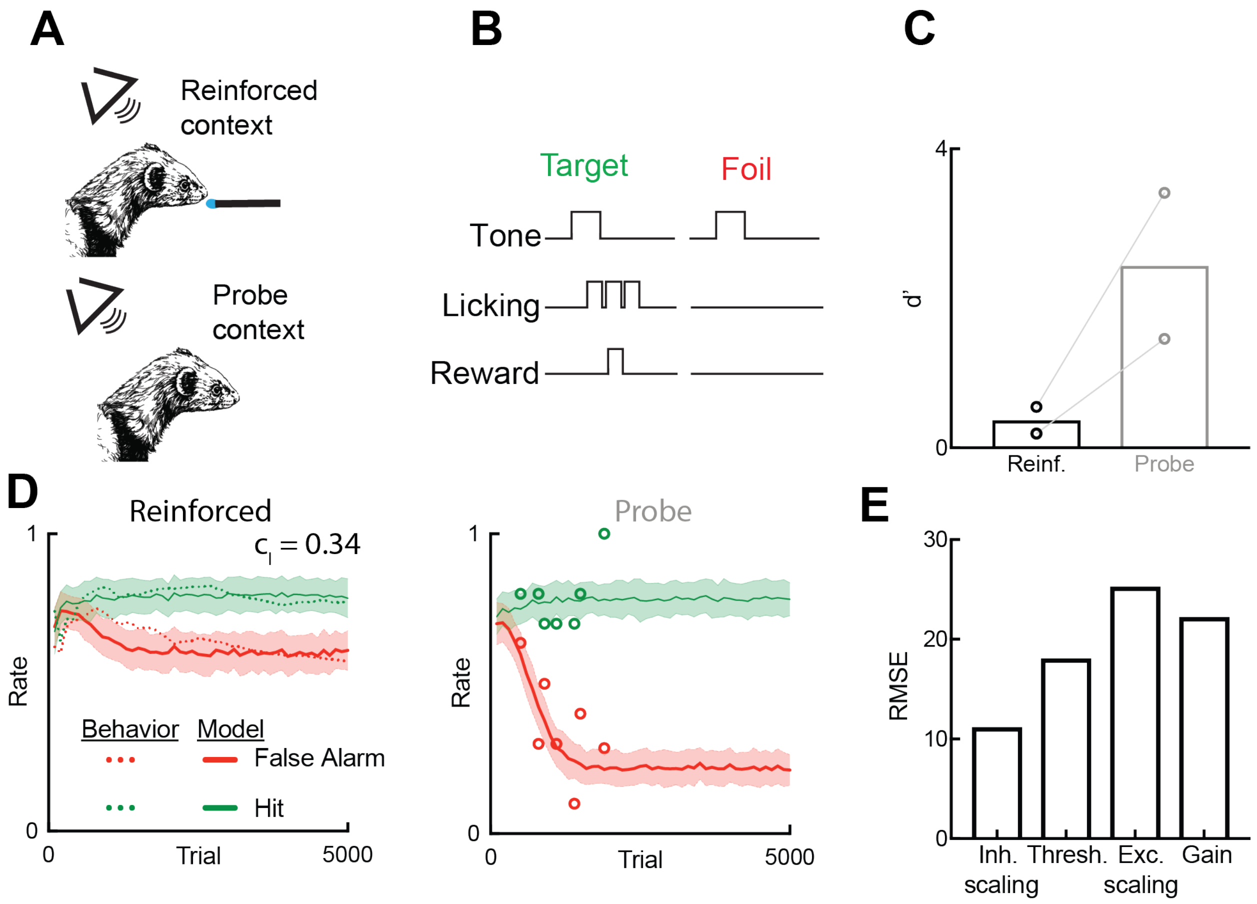


**Figure S2: Dissociation of acquisition and expression in ferrets. A**, Behavioral schematic showing task set-up in the reinforced and probe contexts. **B**, Ferrets were trained to lick to one auditory stimulus for a small water reward, and to withhold licking to another foil auditory stimulus to avoid a short time out. **C**, Behavioral sensitivity (d') of ferrets in the reinforced and probe context in the intermediate stages of training (Reinforced d': 0.37±0.18, mean±s.e.m; Probe d': 2.44±0.98, mean±s.e.m, N=2 ferrets). Both ferrets were tone responsive (i.e., licked selectively following stimulus presentation and not during ITI's), but discriminated poorly in the reinforced context. **D**, Learning trajectory of one ferret in both the reinforced and probe context, along with the performance of inhibitory scaling model ($c_I = 0.303$). **E**, Error rates for each of the four best models tested on the individual ferret whose performance was tracked throughout learning: inhibitory scaling, threshold modulation, excitatory scaling, and gain modulation. Inhibitory scaling performed better than all other tested models across contexts.

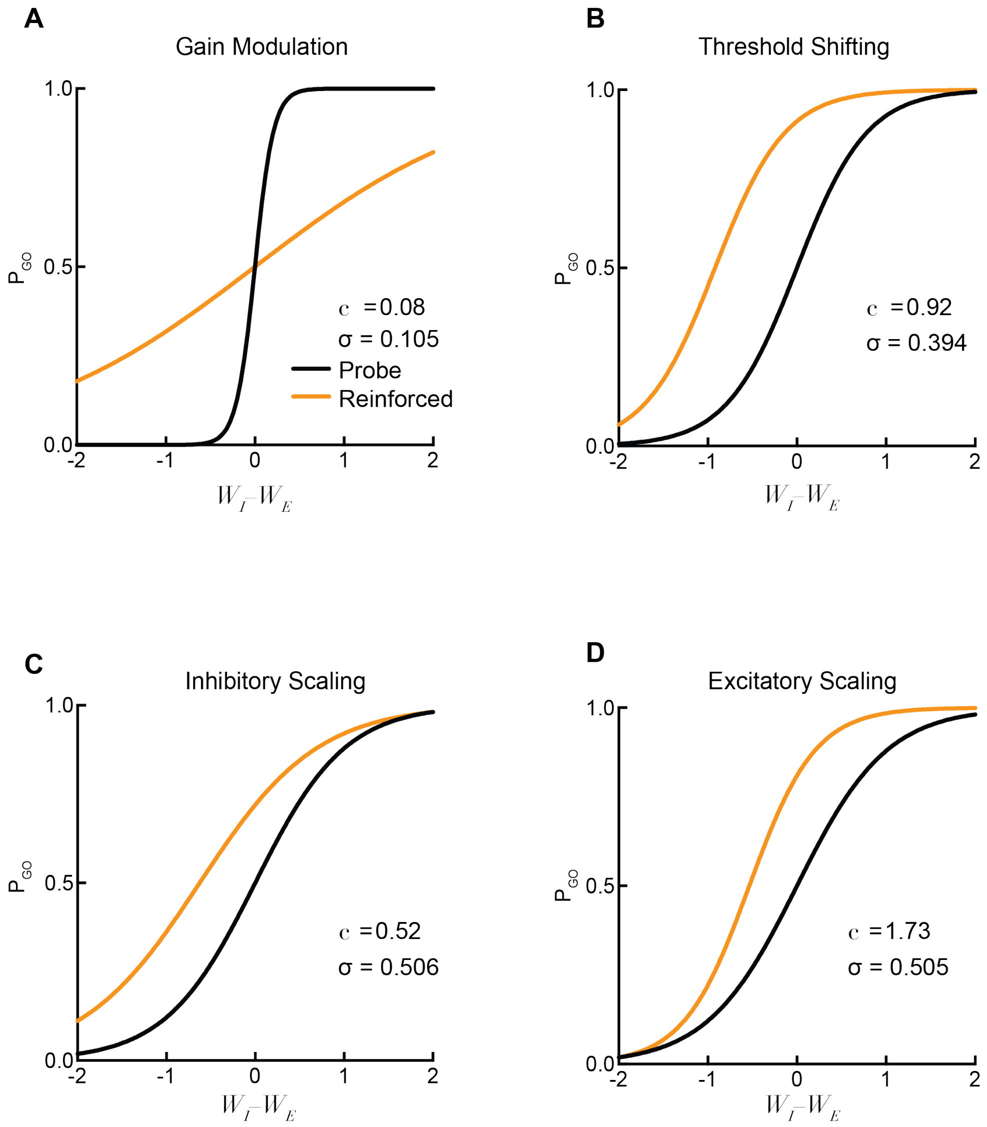


**Figure S3: Scaling factors modulate non-linear decision readouts.** Response probability of the decision-making unit given the excitatory and inhibitory weights, for each of the four models tested: gain modulation (**A**), threshold shifting (**B**), inhibitory scaling (**C**), and excitatory scaling (**D**). In particular, note the flattening of the response probability curve with gain modulation.

\

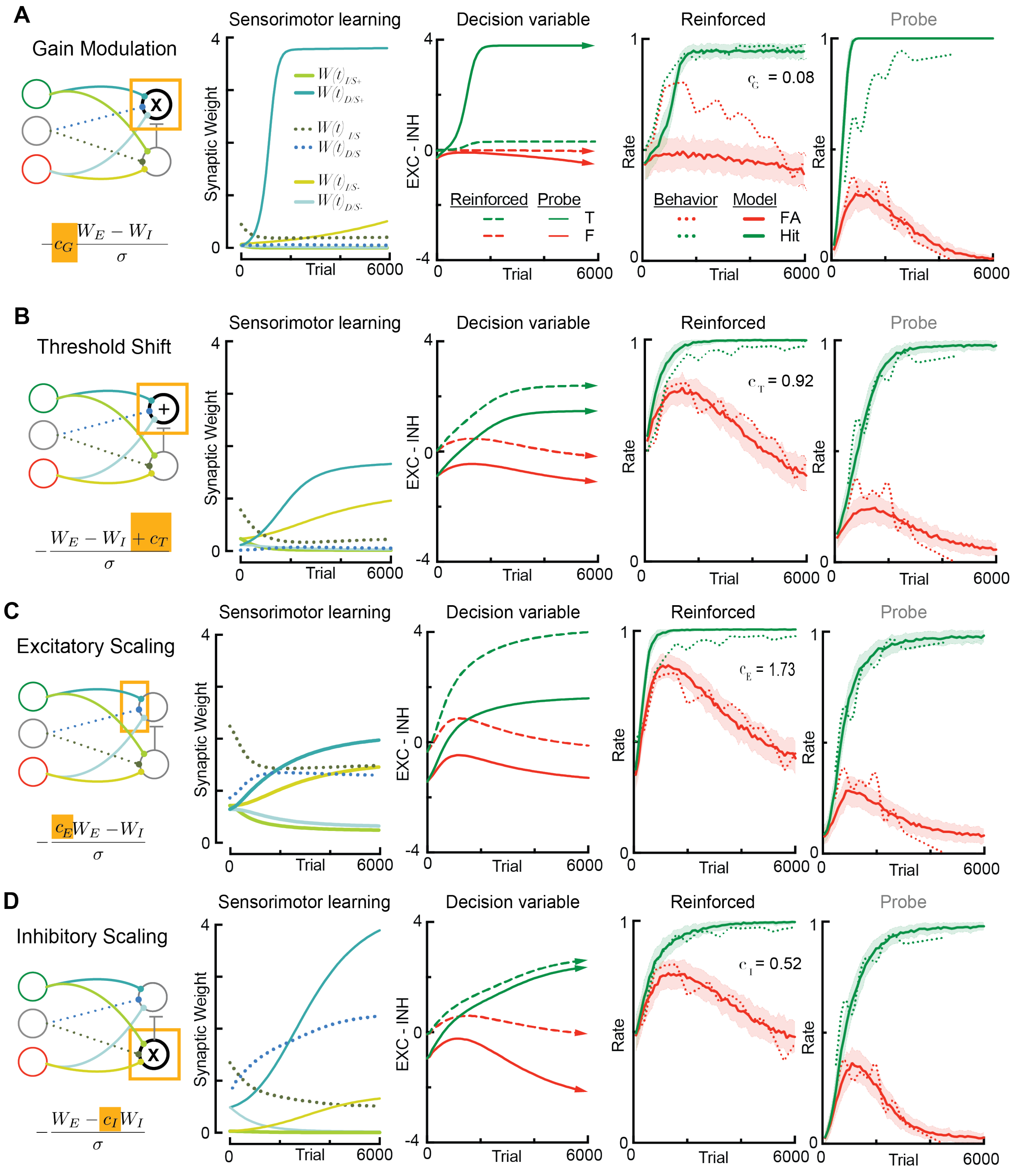


**Figure S4: Non-selective scaling fails to capture parallel learning trajectories. A**, Schematic of multiplicative gain modulation. *Sensorimotor learning*: the synaptic weights over the course of model training using gain-modulation. *Decision variable*: the net input to the decision-making unit for the target (green) and foil (red) tone over the course of learning. Solid lines indicate input during probe trials (baseline model), dashed lines indicate input in the reinforced context (gain modulated, $c_G$=0.08). *Reinforced:* performance of gain model versus average behavioral observations from mice performing the auditory go/no-go task in the reinforced context ($c_G$ =0.08). *Probe context:* performance of gain model versus behavioral observations of 7 mice in the probe context ($c_G$ = 1). **B-D**, All panels are similar to above but the contextual modulation is provided by additive threshold-shifting (b), excitatory scaling (c), or inhibitory scaling (d). Across contexts, the inhibitory scaling model performed significantly better than all other implemented models (**Figures S6f, S7f**).

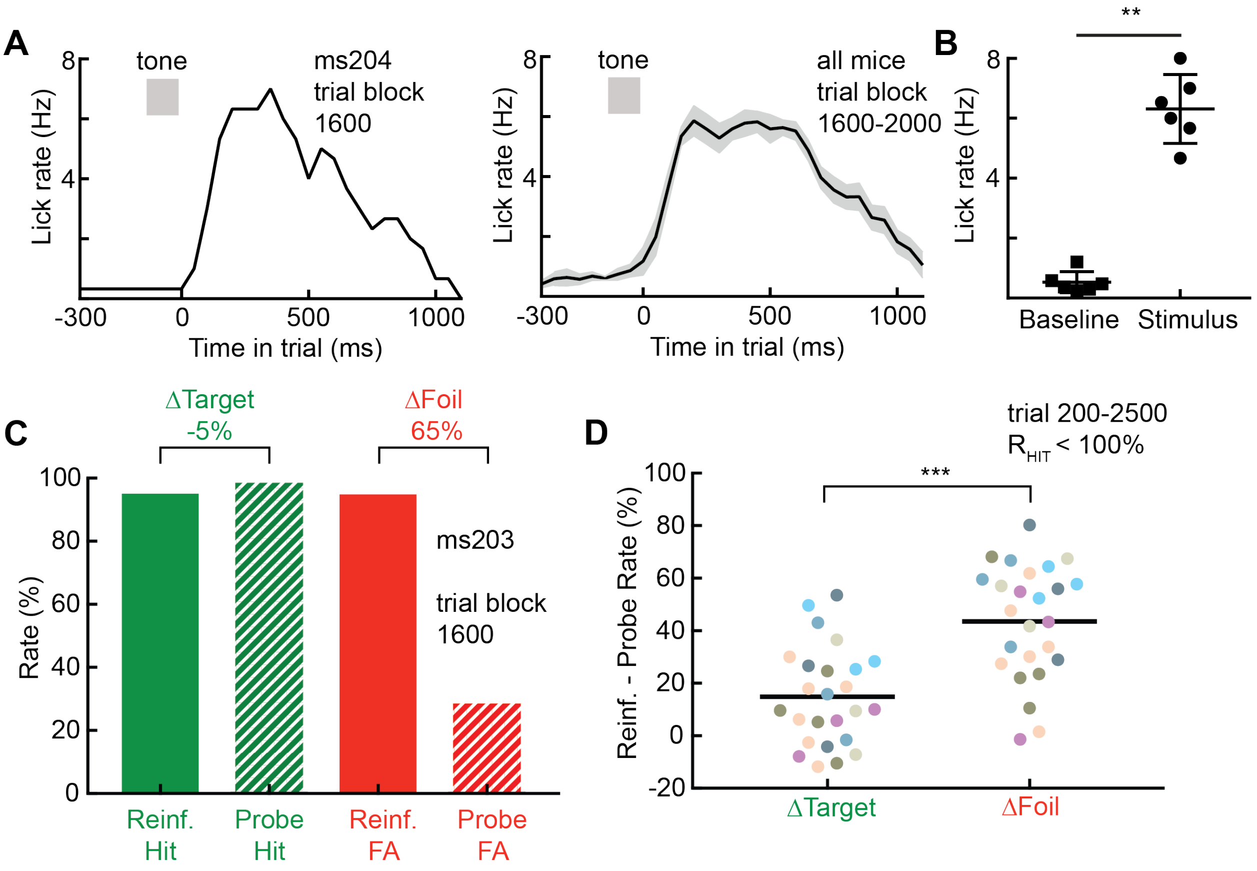


**Figure S5: Compound learning of licktube-reward association does not account for performance differences. A,** Peri-stimulus histogram of lick responses of an example mouse (left) and average of six mice (right). Responses are in the reinforced context and during sessions in which there is a large performance gap between reinforced and probe contexts (i.e., trial block 1500-2000). **B,** Nominal baseline licking in comparison to stimulus evoked responses for trial block 1500-2000 in the reinforced context (N=6 mice, t(5)=9.67, p=0.0002, Student's paired two-tailed t-test **C**, Example of non-linear change in hit and false-alarm rates between the reinforced and probe context: reinforced hit rate: 95.1%, probe hit rate: 100%; reinforced false-alarm rate: 94.9%, probe false-alarm rate: 30%. Note that hit rate in the reinforced context was below 100%. **D**, Difference in hit rate between reinforced and probe context (green) versus difference in false alarm rate between reinforced and probe context (red). Dots indicate individual training blocks (reinforced trials immediately followed by probe trials) where the reinforced hit rate was below 100%, and colors represent individual animals. All training blocks occurred between trials 300 and 2500. ΔTarget rates were significantly lower than ΔFoil rates (ΔTarget: 14.9±18.7%, ΔFoil= 48.6±18.7%, n=25 training blocks, N=7 mice, average Δ for each animal used for statistics, $t(6)=-11.357$, $p=2.79\times10^{-5}$, Student's paired two-tailed t-test).

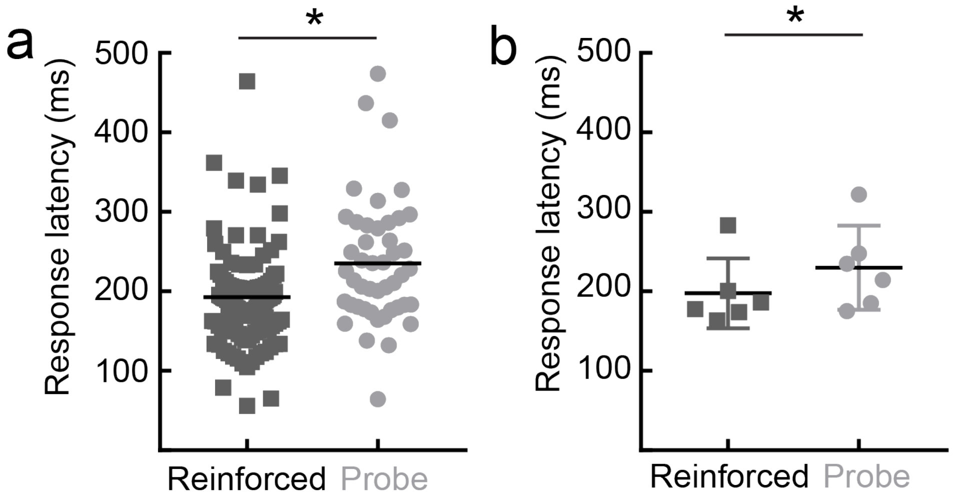


**Figure S6: Response latency is longer during probe trials early in learning. A**, Across mice, response latency (time to first lick following stimulus presentation) is lower in the reinforced (left) context than in the probe (right) context (probe context: 235.4±11.6 ms, mean±s.e.m, n=46 trials; reinforced context: 189.1±5.5 ms, mean±s.e.m., n=132 trials; t(176) = 3.315, p=0.001, unpaired Student's two-tailed two-sample t-test). **B**, Summary of data in **A**, represented as the average response latency of individual mice. Response latency was significantly lower in the reinforced context than in the probe context (probe context: 229.8±21.6 ms, mean±s.e.m, n=46 trials; reinforced context: 197.5±17.9 ms, mean±s.e.m., N=6 mice; t(5) = 3.509, p=0.017, Student's paired two-tailed t-test).

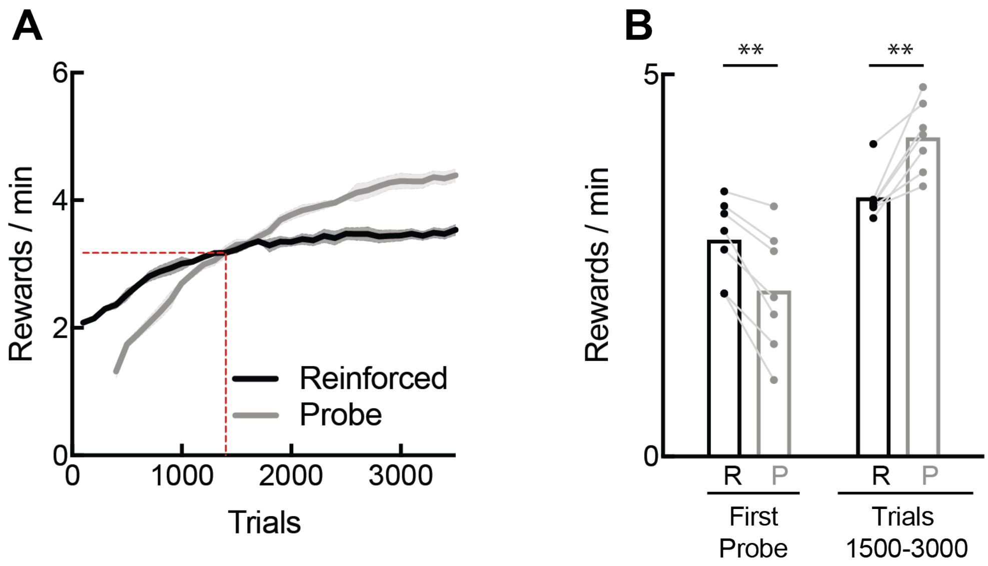


**Figure S7: Exploration in the reinforced context early in learning improves reward rates**. **A**, Average number of reward attained per minute in the reinforced versus probe context for mice the go/no-go task over the course of learning (N = 7 mice). The number of rewards per minute attained in the probe context exceeded the reward rate in the reinforced context after 1400 trials. Reward rates were computed from the average behavioral hit and false alarm rates in blocks of 100 trials. **B**, Reward rates of animals in the reinforced versus probe context at the onset of training versus an intermediate stage of training. During the first probe session, animals attained a lower reward rate in the probe context than in the reinforced context because of higher action rates in the reinforced context session immediately preceding it (reinforced: 2.83±0.20 rewards / min; probe: 2.17 ± 0.30 rewards / min, mean±s.e.m., N = 7 mice; $F(3,18) = 22.44$, $p < 0.01$, one-way repeated measures ANOVA followed by Tukey's post-hoc correction), but with similar discrimination. Across trials 1500-3000, animals attained a higher reward rate in the probe context than in the reinforced context due to improved discriminability (reinforced: 3.39±0.12 rewards / min; probe: 4.18 ± 0.18 rewards / min, mean±s.e.m., N = 7 mice; $F(3,18) = 22.44$, $p < 0.01$, one-way repeated measures ANOVA followed by Tukey's post-hoc correction). Dots indicate individual animals.

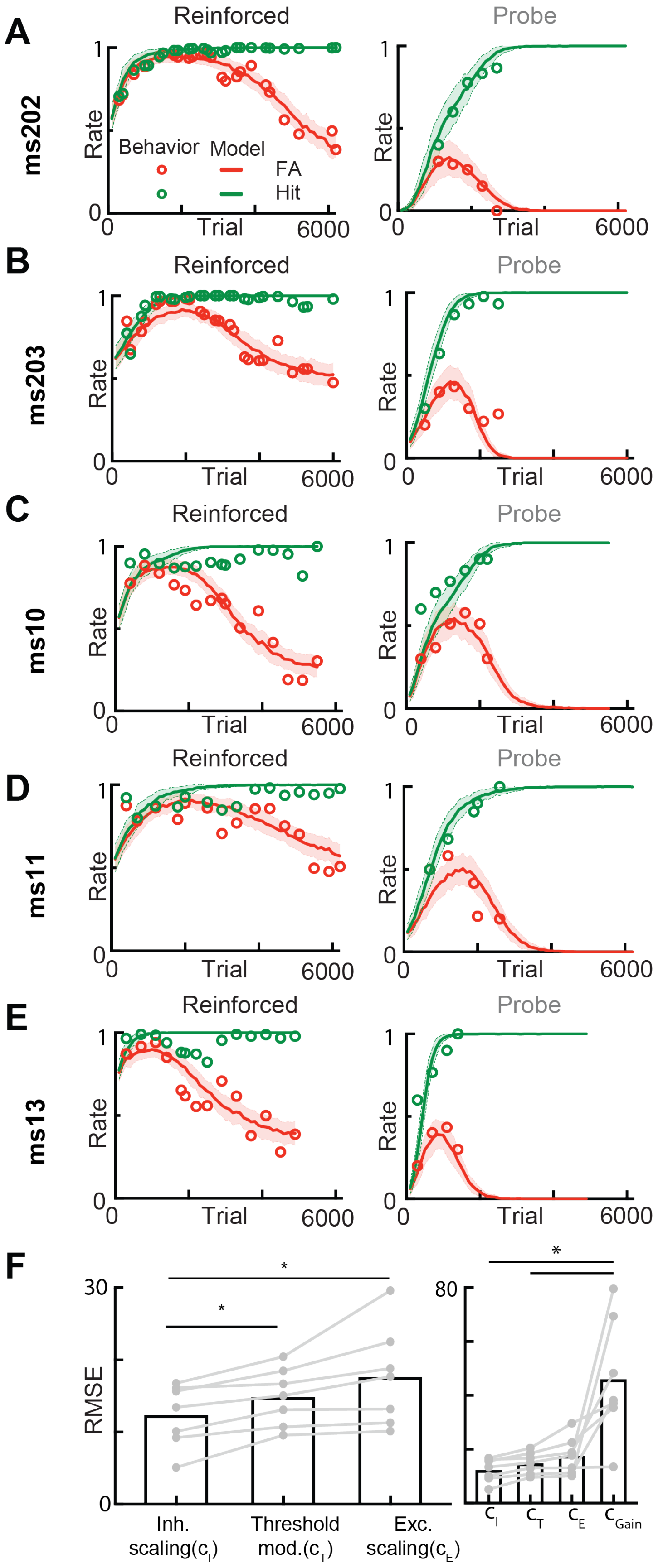


**Figure S8: Inhibitory scaling optimally captures individual variability.** Sample model fits to individual mice for the inhibitory scaling model across both behavioral contexts (**A-E**). **F**, Error rates for each of the three best models tested across individual animals: inhibitory scaling, threshold modulation, and excitatory scaling ($F(3,18) = 1.051$; *, $p<0.05$, repeated-measures one-way ANOVA followed by Tukey's post-hoc correction). Inhibitory scaling performed significantly better than all other tested models across contexts. Right panel similar to left, but different y-axis scale to include error rates for gain model.

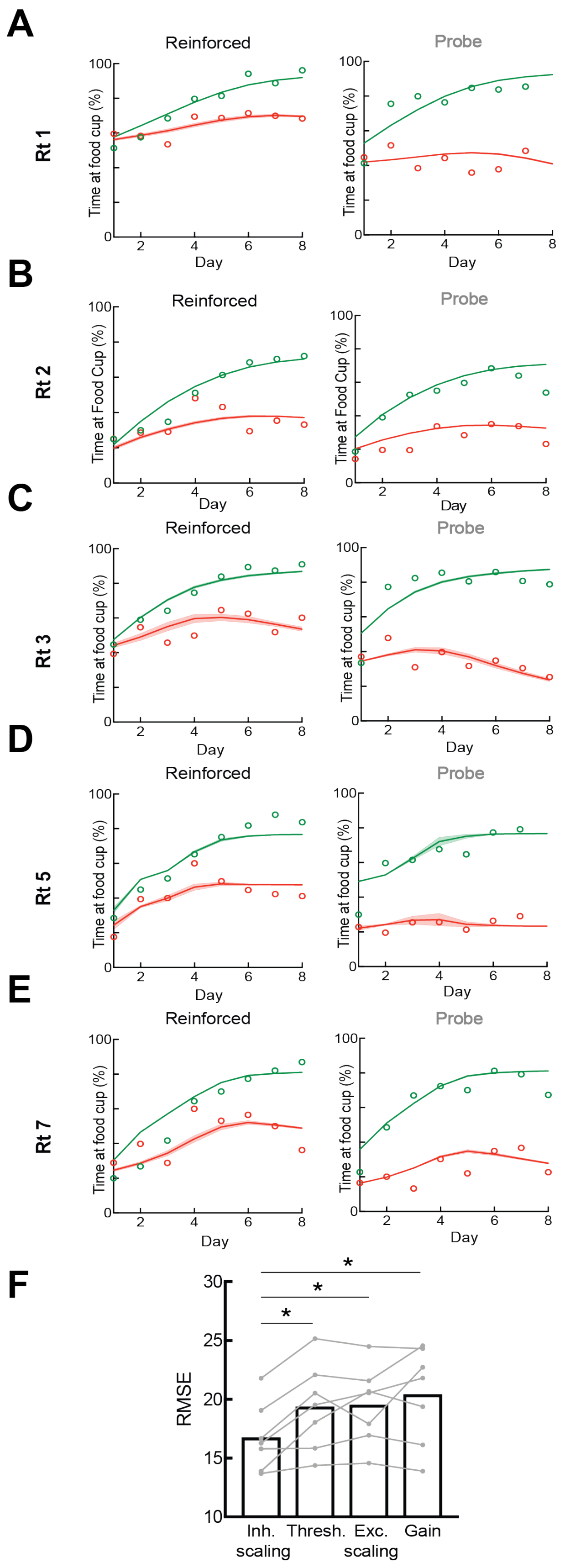


**Figure S9: Inhibitory scaling optimally captures individual variability in freely behaving rats. A-E,** Sample model fits to individual rats learning a Pavlovian serial feature negative discrimination task the inhibitory scaling model across both behavioral contexts. **F**, Error rates for each of the four best models tested across individual animals: inhibitory scaling, threshold modulation, excitatory scaling, and gain modulation ($F(3,18) = 8.829$; *, $p<0.05$, repeated-measures one-way ANOVA followed by Tukey's post-hoc correction). Inhibitory scaling performed significantly better than all other tested models across contexts.

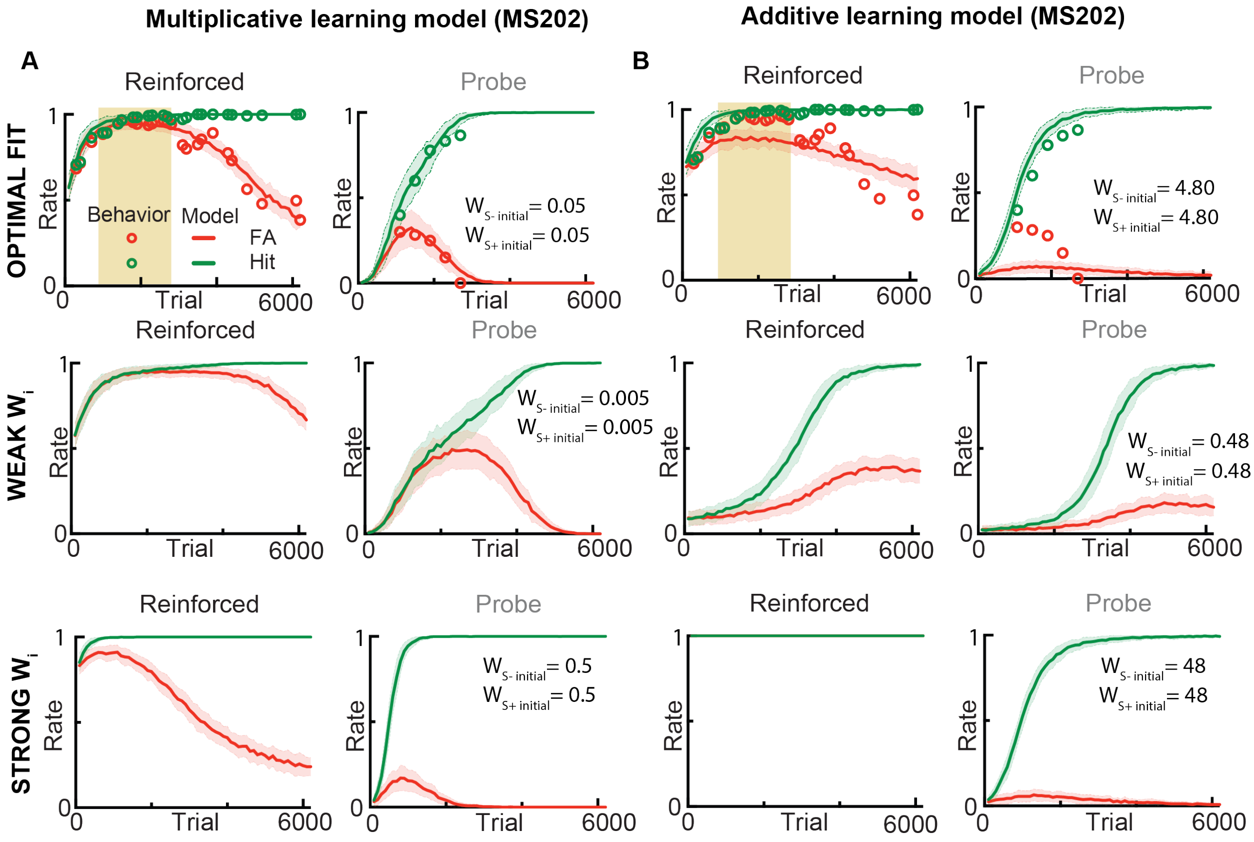


**Figure S10: Multiplicative learning is necessary to capture initial delay in performance improvement in the reinforced context. A**, Top: optimized fit in the reinforced and probe context of a multiplicative model for an individual animal with a strong delay period. Middle: decreasing the initial weights from the tone-selective population (S+/S-) increased the delay period. Behavioral improvement occurred approximately exponentially. Bottom: Increasing the initial weights from the tone-selective population decreased the length of the delay period, making the learning trajectory increasingly sigmoidal in the reinforced context. **B**, Same as **A** but using additive learning rules. Learning trajectories were strongly impacted by initial weight changes of the same magnitude as **A**, but this impact was less restricted to the delay period. An additive model could not account for the learning trajectories of some individual animals (top).
